# Bioorthogonal metabolic labeling of the virulence factor phenolic glycolipid in mycobacteria

**DOI:** 10.1101/2023.11.28.569059

**Authors:** Lindsay E. Guzmán, C. J. Cambier, Tan-Yun Cheng, Kubra F. Naqvi, Michael U. Shiloh, D. Branch Moody, Carolyn R. Bertozzi

## Abstract

Surface lipids on pathogenic mycobacteria modulate infection outcomes by regulating host immune responses. Phenolic glycolipid (PGL) is a host-modulating surface lipid that varies among clinical *Mycobacterium tuberculosis* strains. PGL is also found in *Mycobacterium marinum* where it promotes infection of zebrafish through effects on the innate immune system. Given the important role this lipid plays in the host-pathogen relationship, tools for profiling its abundance, spatial distribution, and dynamics are needed. Here we report a strategy for imaging PGL in live mycobacteria using bioorthogonal metabolic labeling. We functionalized the PGL precursor *p*-hydroxybenzoic acid (*p*HB) with an azide group (3-azido *p*HB). When fed to mycobacteria, 3-azido *p*HB was incorporated into the cell surface, which could then be visualized via bioorthogonal conjugation of a fluorescent probe. We confirmed that 3-azido *p*HB incorporates into PGL using mass spectrometry methods and demonstrated selectivity for PGL-producing *Mycobacterium marinum* and *Mycobacterium tuberculosis* strains. Finally, we applied this metabolic labeling strategy to study the dynamics of PGL within the mycobacterial membrane. This new tool enables visualization of PGL which may facilitate studies of mycobacterial pathogenesis.

## Introduction

*Mycobacterium tuberculosis (M. tb)*, the pathogen responsible for tuberculosis (TB), remains the leading cause of death from a bacterium.^1^ A factor that contributes to *M. tb*’s success is its unique lipid-rich cell envelope (Figure 1a).^2^ Many mycobacterial cell-surface lipids play important roles in virulence by modulating the host immune system.^3^ Two structurally related virulence lipids are phthiocerol dimycocerosate (PDIM) and phenolic glycolipid (PGL) (Figure 1b), which are found in the outermost layer of the mycomembrane.^4^ PDIMs and PGLs contain a lipid core with two esterified mycocerosic acid side chains. PGLs are heterogenous with respect to their lipid chain lengths and functionalization with methoxy, hydroxy, or keto groups.^4^ Additionally, PGLs have a phenol moiety and a species-dependent glycan. *Mycobacterium marinum* (*M. marinum*) also produces PGL and is required for virulence.^5^ However, the *M. marinum* PGL glycan structure differs from those found in *M. tb* (Figure 1c).^6^ Each component of the PGL structure contributes to its effects on virulence.^7^

**Figure 1.**
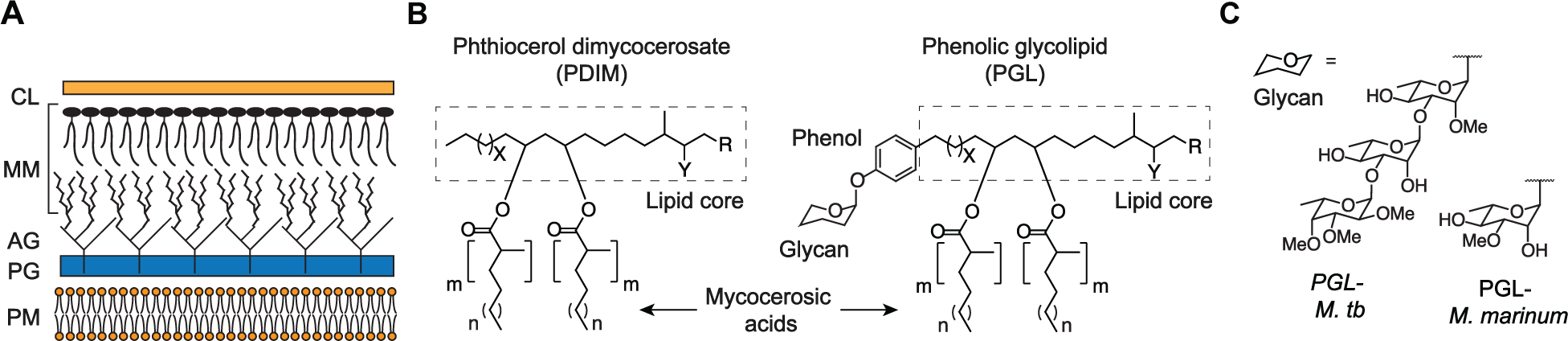
Phenolic glycolipid (PGL) is a mycobacterial cell-surface virulence factor. (A) Layers of the mycobacterial cell wall. CL = capsular layer, MM = mycomembrane, AG = arabinogalactan, PG = peptidoglycan, and PM = plasma membrane. (B) Simplified representative chemical structures of PDIM and PGL. PDIM and PGL structures are heterogeneous, and reports vary in literature. For *M. tb*: X = 14-16; Y = methoxy, keto, or hydroxy; m = 3-5; n = 15-17; and R = CH_3_ or H. For *M. marinum*: X = 14-16; Y = methoxy, keto, or hydroxy; m = 3-4; n = 16-18; and R = CH_3_. (C) Glycans of PGL vary according to the species of mycobacteria.

The immunomodulatory effects of PGL are dependent on the species. *M. tb* PGLs are found in hypervirulent Lineage 2 strains, such as HN878.^8–9^ These PGLs have been shown to suppress the secretion of proinflammatory cytokines TNF-α, IL-6, and MCP-1.^7–8, 10^ In the zebrafish infection model, *M. marinum* PGLs allow bacterial transfer to permissive monocytes^11^ through production of CCL2.^12^ Overall, PGLs are important virulence factors that give rise to immunomodulatory host responses.^13–14^

The ability to image PGL could be empowering for studies of its distribution and dynamics on mycobacterial cells. Unlike proteins that can be engineered for biosynthesis with fluorescent protein labels, lipids require chemical tools for labeling and visualization. One such approach is to modify lipids with bioorthogonal handles (e.g. azides or alkynes), then conjugate them to fluorescent probes in living systems.^15–16^ We and others have used metabolic labeling to incorporate a bioorthogonal handle into trehalose monomycolate (TMM), a major immunogenic lipid of the mycobacterial cell envelope.^17–19^ Additionally, we developed a chemical approach to visualize a fluorescent PDIM during infection of zebrafish with *M. marinum*.^20^

Here, we report a metabolic labeling strategy to image PGL on live mycobacterial cells. We synthesized an azide-functionalized PGL precursor that is incorporated into native PGL within the outer membrane of the model mycobacterial species, *M*. *marinum*. We characterized azide-modified PGL (PGL-N_3_) using mass spectrometry, demonstrated selectivity of labeling in *M. marinum* and *M. tb*, and used this imaging tool to study PGL dynamics within the mycobacterial membrane. The ability to image PGL on live mycobacteria adds to the toolkit for experimental studies of mycobacterial lipid biology.

## Results and Discussion

The biosynthesis of *M. tb* PGLs is a complex multistep process (Scheme 1).^21^ It is hypothesized that PGL biosynthesis is conserved across PGL-producing mycobacterial species^22^ apart from the glycan. The first committed step in PGL biosynthesis is loading of *p*-hydroxy benzoic acid (*p*HB) onto the fatty-acid-CoA ligase, FadD22,^23^ and elaboration by the type 1 polyketide synthase pks15/1.^24–26^ A variety of polyketide synthases^27^ and other enzymes^28^ further extend and embelish the lipid core and decorate it with methoxy, keto, or hydroxy functional groups to form phenolphthiocerol, phenolphthiodiolone, or phenolphthiotriol lipid cores.^29–30^ In parallel, mycocerosic acids are synthesized by the mycocerosic acid synthase (mas)^31^ then condensed with the lipid core by the enzyme PapA5 to form *p*-hydroxyphenol PDIM.^32^ The glycan of PGL is then decorated by several glycosyl and methyltransferases.^29, 33–34^ Finally, PGL is shuttled to the cell surface by the lipid transporter Mmpl7 and its auxiliary proteins.^35^

**Scheme 1.**
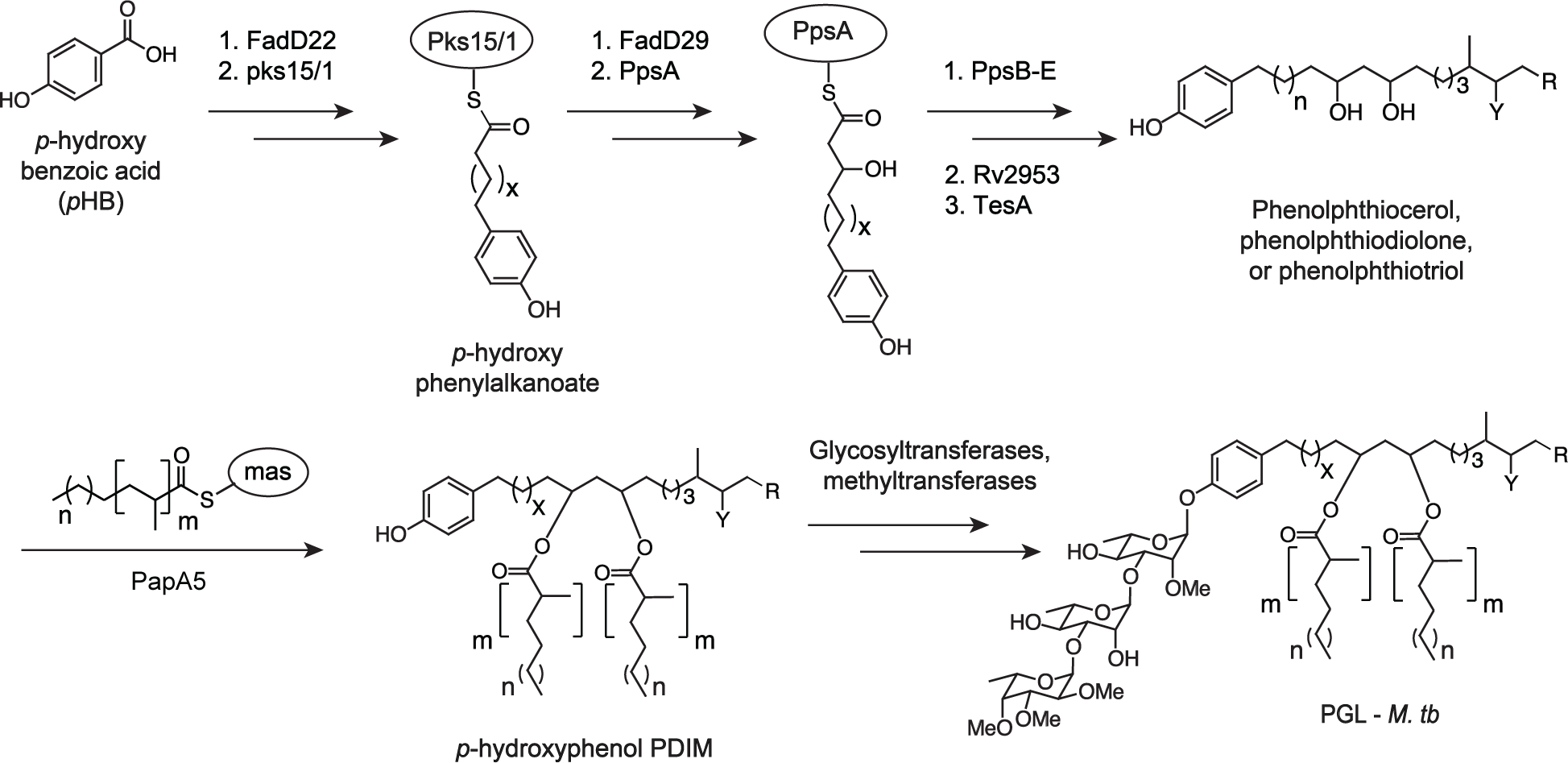
Simplified PGL biosynthetic pathway for *M. tb*. PGL structures are heterogeneous, and reports vary in literature, but typically X = 14-16; Y = methoxy (phenolphthiocerol), keto (phenolphthiodiolone), or hydroxy (phenolphthiotriol); m = 3-5; n = 15-17; and R = CH_3_ or H.

We focused on *p*HB as a key intermediate that could be modified with an azide group. Previous work has shown that exogenous radiolabeled *p*HB is taken up by mycobacterial cells and metabolically incorporated into cell-surface PGL.^8^ While *p*HB is also used in the biosynthesis of other *p*-hydroxybenzoic acid derivatives (*p*HBADs),^36–39^ these metabolites are not associated with the cell envelope — they are either cytosolic or secreted — and therefore would not be expected to confound the visualization of membrane-associated PGL. We synthesized both 2- and 3-azido *p*HB as described in the supporting information. We tested these derivatives as substrates for metabolic labeling of cell surface PGL using the workflow shown in Figure 2a. We used *M. marinum* as a model mycobacterial species based on its close genetic relationship to *M. tb* and its prior use in studies of mycobacterial pathogenesis.^40–41^

**Figure 2.**
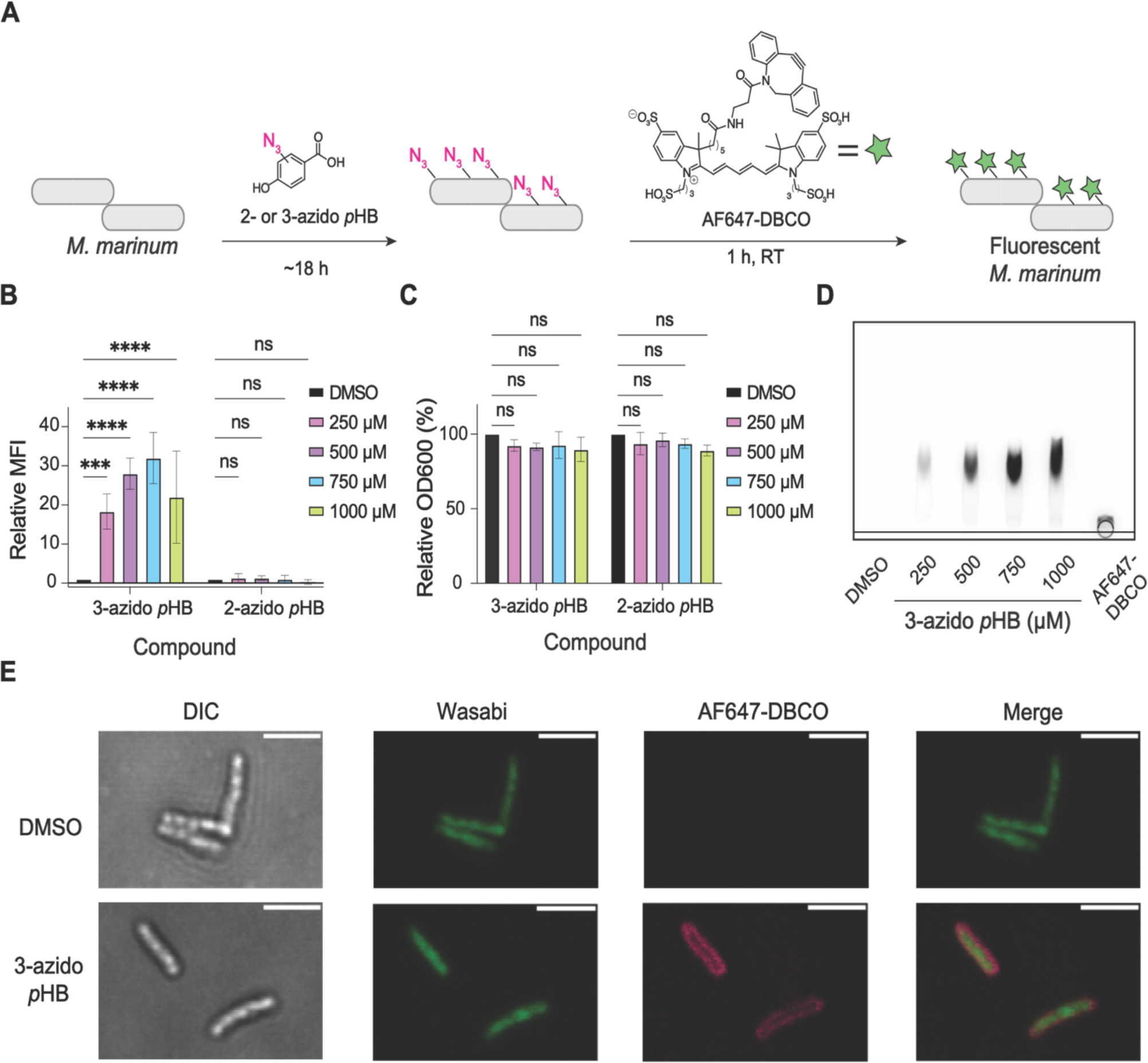
Fluorescent labeling of *M. marinum* upon treatment with 3-azido *p*HB. (A) Workflow of labeling experiments with 2- or 3-azido *p*HB. *M. marinum* were treated for 18 h in the presence of various concentrations of 2- or 3-azido *p*HB followed by staining with DBCO-AF647 (30 μM). Cells were then fixed and analyzed by flow cytometry. (B) Flow cytometry analysis of labeled cells. Relative MFI is determined by normalizing against DMSO control. Flow cytometry data are the average of biological triplicates. Statistical analysis was performed using a two-way ANOVA followed by a Dunnett’s multiple comparisons test. Significance is represented as follows: \*\*\**p* < 0.001, \*\*\*\**p* < 0.0001, and ns (not significant) for *p* > 0.05. (C) Effects of bacterial growth (OD600) of *M. marinum* treated with 2- or 3-azido *p*HB. Statistical analysis was performed using an ordinary two-way ANOVA followed by Šídák’s multiple comparisons test. Significance is represented where ns (not significant) for *p* > 0.05. (D) Thin layer chromatography (TLC) analysis of crude lipid extracts from *M. marinum* cells treated with 3-azido *p*HB and stained with AF647-DBCO. Crude lipid extracts (50 µg) or AF647-DBCO (1 µg) were loaded onto a silica gel 60 TLC plate, which was then developed with 4:6 methanol:chloroform. (E) Confocal images of *M. marinum* cells treated with 750 µM 3-azido *p*HB and stained with AF647-DBCO. Scale bar = 2 µm.

*M. marinum* cells were treated with various concentrations of 2- or 3-azido *p*HB (250 – 1000 µM) for 18 h. The bacteria were then washed, stained with the fluorophore AF647-dibenzocyclooctyne (AF647-DBCO), fixed with 4% paraformaldehyde/2.5% glutaraldehyde, and analyzed by flow cytometry (Figure 2b). When bacteria were treated with 2-azido *p*HB, there was no increase in mean fluorescence intensity (MFI) in reference to the DMSO control. However, when *M. marinum* were treated with 3-azido *p*HB, we saw a significant and dose-dependent increase in MFI up to 750 µM with modestly reduced fluorescence at the highest dose (Figure 2b). We also assessed the effects of 2- or 3-azido *p*HB treatment on PGL production qualitatively by thin layer chromatography (TLC). When lipids from 2-azido *p*HB-treated cells were extracted with chloroform/methanol and analyzed by TLC, we observed a significant decrease in PGL’s abundance at all concentrations tested (Figure S1). This outcome suggests that 2-azido *p*HB strongly inhibits PGL biosynthesis. By contrast, treatment of cells with 3-azido *p*HB qualitatively showed much smaller reductions in PGL production by TLC, which were apparent at the highest concentrations tested (Figure S1). Neither 2- nor 3-azido *p*HB significantly affected cell growth as measured by OD600 (Figure 2c). Importantly, no fluorescence labeling was observed when cells were treated with natural *p*HB followed by the labeling reagent, AF647-DBCO, indicating that an azide is required for increases in MFI (Figure S2). Given the outcome of these experiments, we focused on using 3-azido *p*HB as a metabolic label for the remainder of this study.

To determine whether fluorescence labeling resulted from incorporation of 3-azido *p*HB into cell surface lipids, we treated *M. marinum* with 3-azido *p*HB followed by AF647-DBCO, extracted lipids with chloroform/methanol, and separated them by TLC (Figure 2d). The extracts contained one major fluorescent species with an R_f_ consistent with a lipid that is modified with a charged fluorescent dye.

We then used confocal microscopy to determine localization of fluorescence in labeled *M. marinum* (Figure 2e). *M. marinum* expressing a green fluorescent protein (wasabi) were treated with 750 µM 3-azido *p*HB followed by staining with AF647-DBCO. As shown in Figure 2e, AF647 fluorescence was only observed on the cells treated with 3-azido *p*HB, consistent with our flow cytometry data (Figure 2b). The fluorescence was concentrated on the outer membrane of the cells, where PGL is located. No difference in cell morphology was observed in comparison to the DMSO control, suggesting that incorporation of azides into PGL has no dominant effect on *M. marinum* cellular macrostructures.

To confirm that 3-azido *p*HB was incorporated into PGL, we used high-performance liquid chromatography quadrupole time-of-flight mass spectrometry (HPLC-Q-TOF-MS) to analyze *M. marinum* lipid extracts. The sample without 3-azido *p*HB treatment served as a control. Endogenous PGL was detected as an ammonium adduct (1523.3928 *m/z*) (Figure 3a). We generated a table of predicted molecular formulas and *m/z* values of the ammonium adducts for known PGL lipoforms (Supplemental Table 1). Because PGL-N_3_ would have a net mass gain of 41.0014 *m/z* due to replacement of a proton by an azide group, we calculated the theoretical *m/z* values for PGL-N_3_ lipoforms (Supplemental Table 1).

**Figure 3.**
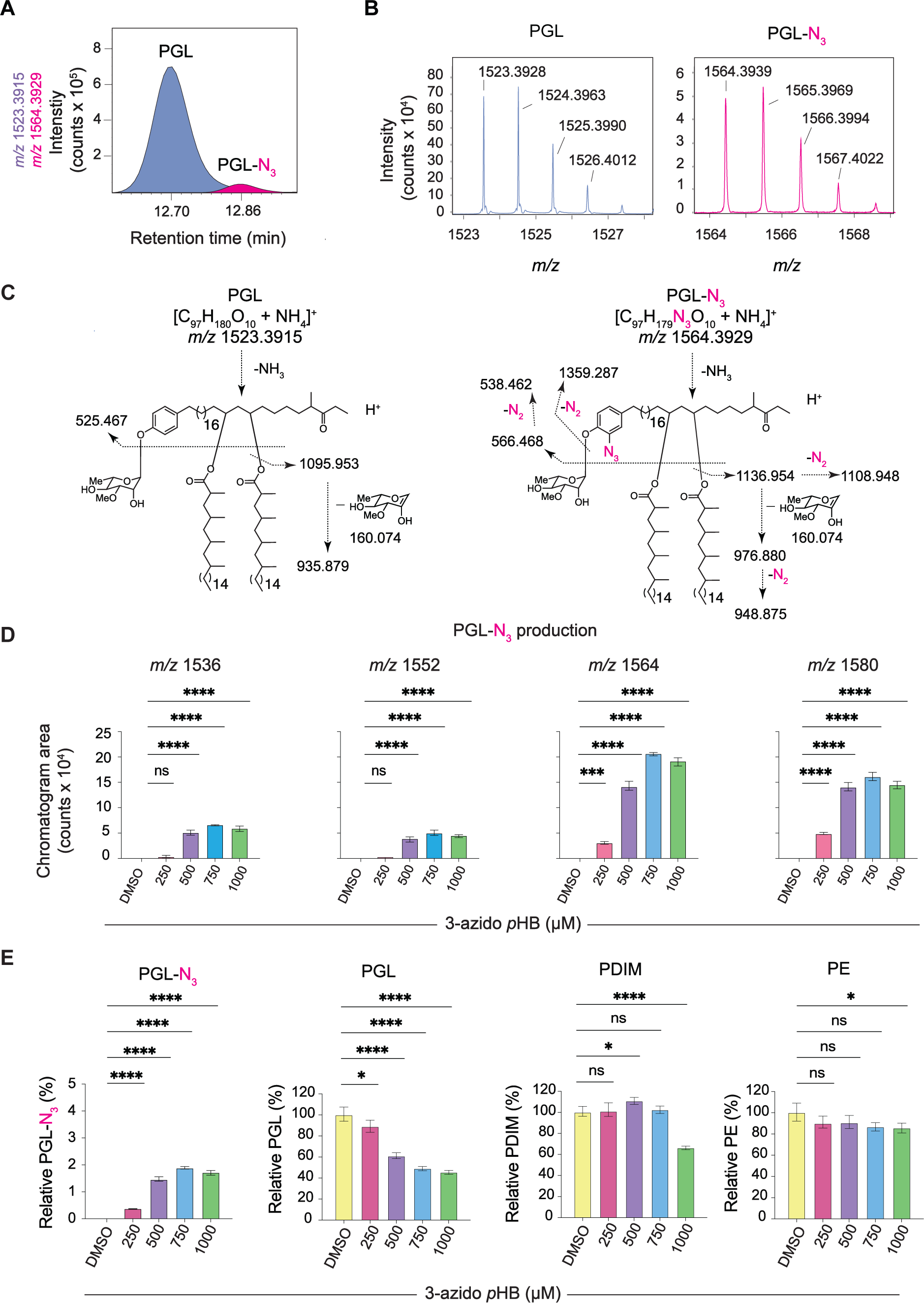
Mass spectrometry (MS) analysis of crude lipid extracts from 3-azido *p*HB-treated *M. marinum*. (A) The ion chromatograms of the representative species of PGL (1523.3915 *m/z*) and PGL-N_3_ (1564.3929 *m/z*) detected in the total lipid extracts of *M. marinum* treated with 3-azido *p*HB (750 µM) were generated by positive-mode, reversed-phase HPLC-Q-ToF-MS. (B) Mass spectra of indicated PGL and PGL-N_3_ species. (C) Collision-induced MS (CID-MS) of PGL and PGL-N_3_ showing diagnostic fragments corresponding to loss of one mycocerosic acyl moiety (PGL: 1095.953 *m/z*; PGL-N_3_: 1108.948 *m/z* with spontaneous loss of N_2_ from the N_3_ group), loss of one mycocerosyl moiety plus the monosaccharide (PGL: 935.879 *m/z*; PGL-N_3_: 976.880 *m/z* and 948.866 *m/z* with spontaneous loss of N_2_ from the N_3_ group), and loss of both mycocerosic acyl moieties plus the monosaccharide (PGL: 525.467 *m/z*; PGL-N_3_: 566.468 *m/z* and 538.462 *m/z* with spontaneous loss of N_2_ from the N_3_ group). (D) Effects of *M. marinum* treated with 3-azido *p*HB on the abundance of PGL-N_3_ species as quantified by MS. Data represented as three technical replicates. Statistical analysis was performed using a two-way ANOVA followed by a Dunnett’s multiple comparisons test. Significance is represented where \*\*\*\**p* < 0.0001 and ns (not significant) for *p* > 0.05. (E) Effects of *M. marinum* treated with 3-azido *p*HB on abundance of PGL-N_3_, total PGL, PDIM and PE as determined by MS analysis. Quantified mass spectrometry data are representative of three technical replicates. Statistical analysis was performed using a two-way ANOVA followed by a Dunnett’s multiple comparisons test. Significance is represented as follows: \**p* < 0.05, \*\**p* < 0.01, \*\*\*\**p* < 0.0001, and ns (not significant) for *p* > 0.05.

Next, we searched for PGL-N_3_ in the lipid extracts from 3-azido *p*HB-treated cells and found seven ions (1536.3652, 1552.4034, 1564.3939, 1580.4244, 1592.4350, 1606.4491, and 1608.4519 *m/z*) that corresponded to PGL-N_3_ species within a mass error of 10 parts per million (ppm). The mass intervals between these ions corresponded to differences in methoxy or keto groups and chain length variants (Supplemental Table 1). Out of the seven PGL-N_3_ ions identified, four were in major abundance with high mass accuracy (within 5 ppm). One major PGL-N_3_ species (1564.3929 *m/z*) separated from its endogenous PGL counterpart by HPLC (Figure 3a). The identification of PGL-N_3_ in the mass spectrum (Figure 3b) was further validated by collision-induced dissociation mass spectrometry (CID-MS). Like the natural PGL, the PGL-N_3_ species molecule readily lost both mycocerosic acids and the rhamnose glycan in the MS2 spectrum. This yielded lipid core fragments with or without spontaneous loss of N_2_ (Figure 3c), commensurate with literature reports on aryl azide ionization.^42^ These data confirm that 3-azido *p*HB was metabolically incorporated into PGL.

We next sought to quantify the abundance of PGL-N_3_ produced in response to treatment with various concentrations of 3-azido *p*HB. As observed with fluorescence labeling (Figure 2b), a treatment dose of 750 µM gave the maximum MFI by flow cytometry. We hypothesized that fluorescence observed by flow cytometry is due abundance of PGL-N_3_. To address this hypothesis, we measured the effect of 3-azido *p*HB on total production of PGL and PGL-N_3_ by mass spectrometry. The highest amount of PGL-N_3_ detected was 2% of total PGLs at 750 µM 3-azido *p*HB (Figure 3e), which confirms our hypothesis that fluorescence and PGL-N_3_ abundance are correlated. Total PGL abundance was inhibited by 3-azido *p*HB in a dose-dependent manner (∼50% at 1 mM). Additionally, we measured the abundance of other non-PGL lipids. We observed smaller effects on the abundance of PDIM, which shares some biosynthetic steps with PGL (Figure 3e). PDIM levels were unaffected by 3-azido *p*HB at concentrations below 1 mM. Phosphatidylethanolamine (PE), a lipid that does not share any biosynthetic steps with PGL, was mostly unaffected by 3-azido *p*HB treatment except for a slight decrease in abundance at the highest concentration of 1 mM (Figure 3e). Thus, PGL-N_3_ biosynthesis was optimal at a labeling concentration of 750 μM 3-azido *p*HB indicated by both fluorescence and mass spectrometry.

As further confirmation that 3-azido *p*HB primarily labels PGL we performed similar experiments using PGL-deficient mycobacteria. We tested *Mycobacterium smegmatis* (*M. smegmatis*), a commonly used model organism which naturally lacks PGL,^43^ and *M. marinum* mutants that are deficient either in pks15/1 or MmpL7.^11^ PGL deficiency was confirmed by TLC analysis of lipid extracts from these mycobacteria (Figure S3). Indeed, when these mycobacteria were treated with 3-azido *p*HB followed by AF647-DBCO, no significant fluorescence labeling was observed by flow cytometry (Figure 4a). No fluorescent species were detected by TLC analysis of lipid extracts from *M. marinum* mutants treated with 3-azido *p*HB and AF647-DBCO (Figure 4b). Therefore, 3-azido *p*HB appears to exclusively label PGL within the lipid extract.

**Figure 4.**
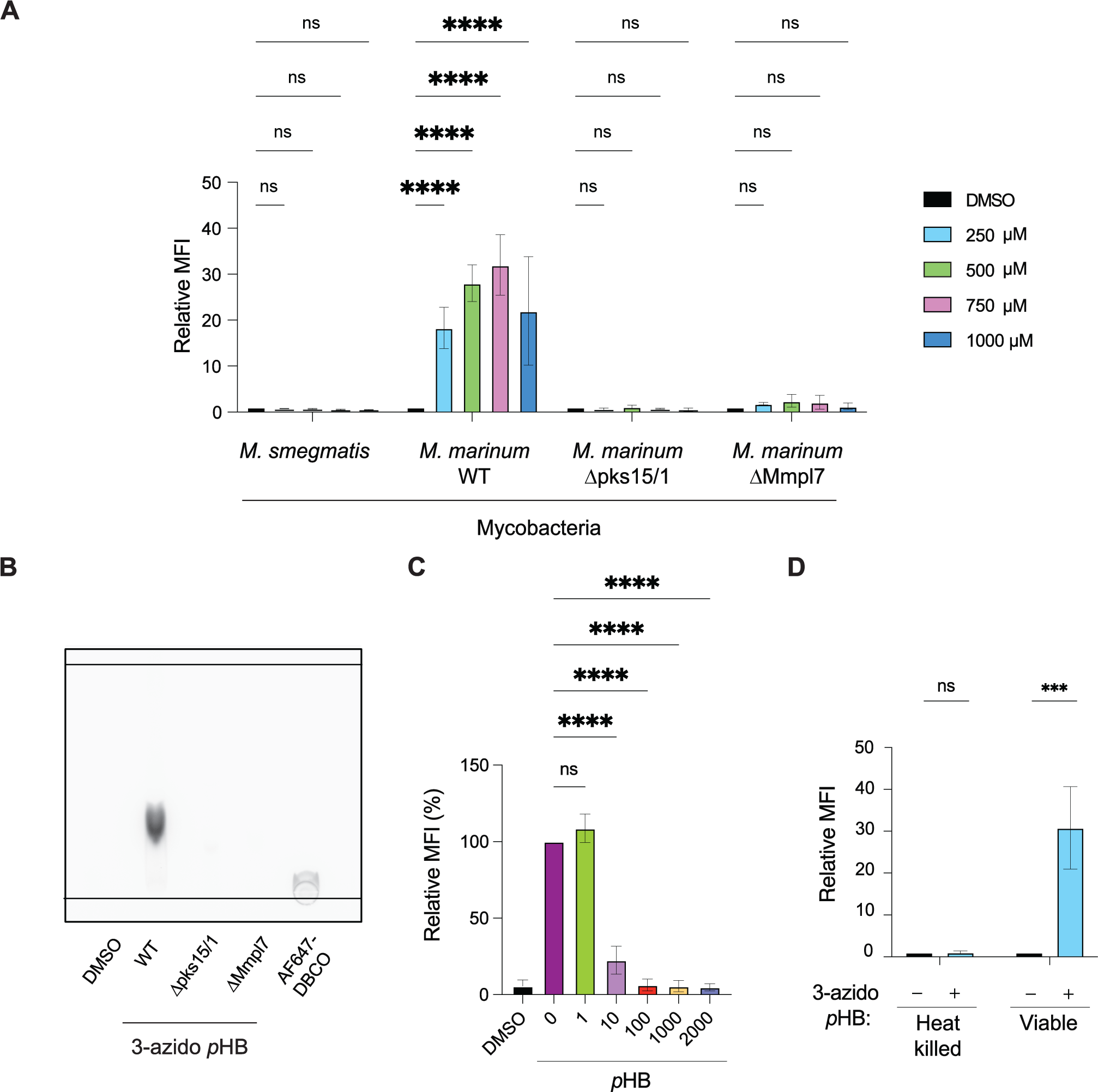
Metabolic incorporation of 3-azido *p*HB only occurs in live *M. marinum* with an intact PGL biosynthetic pathway. (A) Mycobacteria that lack the PGL biosynthetic pathway were treated with various concentrations of 3-azido *p*HB and analyzed by flow cytometry. Flow cytometry data are averages of three biological triplicates. Statistical analysis was performed using an ordinary two-way ANOVA followed by Šídák’s multiple comparisons test. Significance is represented where \*\*\*\**p* < 0.0001 and ns (not significant) for *p* > 0.05. (B) Lipid extracts of PGL-deficient *M. marinum* strains treated with 3-azido *p*HB, stained with AF647-DBCO, and analyzed by TLC. Crude lipid extracts (100 µg) or AF647-DBCO (20 µg) were loaded onto a silica gel 60 TLC plate, which was then developed with 4:6 methanol:chloroform. (C) Competition experiment using various concentrations of *p*HB added to *M. marinum* treated with 750 µM 3-azido *p*HB. Flow cytometry data are averages of three biological triplicates. Relative MFI is determined by normalizing against DMSO control. Statistical analysis was performed using a two-way ANOVA followed by a Dunnett’s multiple comparisons test. Significance is represented where \*\*\*\**p* < 0.0001 and ns (not significant) for *p* > 0.05. (D) *M. marinum* were heat killed (80 °C, 30 min), treated with 3-azido *p*HB for 18 h, stained with AF647-DBCO, and analyzed by flow cytometry. Statistical analysis was performed using a two-way ANOVA followed by a Dunnett’s multiple comparisons test. Significance is represented where *p* < 0.001 and ns (not significant) for *p* > 0.05.

To further test selectivity of 3-azido *p*HB labeling, we performed a competition experiment using natural *p*HB. We treated *M. marinum* with various concentrations of *p*HB in combination with 750 µM 3-azido *p*HB (Figure 4c). As the concentration of *p*HB increased, the fluorescence intensity as measured by flow cytometry decreased in a dose-dependent manner, with complete suppression of metabolic labeling at 100 μM *p*HB. Additionally, we heat-killed *M. marinum* at 80 °C for 30 minutes, which abrogated labeling (Figure 4d). Thus, metabolic labeling of PGL with 3-azido *p*HB only occurs in live cells with active metabolism.

Having demonstrated the ability to image PGL in model organisms, we shifted our attention to virulent *M. tb*. We treated a PGL-producing strain of *M. tb*, HN878, with various concentrations of 3-azido *p*HB and found a significant increase in MFI in comparison to DMSO control (Figure 5a). We noticed that bacterial growth was significantly affected by 3-azido *p*HB treatment (Figure 5b). Next, a PGL-deficient mutant *M. tb* strain, HN878 Δ*pks15/1*, and the naturally PGL-deficient Erdman strain, were treated with 3-azido *p*HB followed by AF647-DBCO (Figure 5c). The absence of PGL in the *M. tb* HN878 Δ*pks15/1* and Erdman strains was confirmed by TLC analysis (Figure S4). We observed no labeling of the two PGL-deficient *M. tb* strains (Figure 5c) which matches observations with PGL-deficient *M. marinum* mutants, confirming the high specificity of the reagent for the PGL pathway (Figure 4a). From these experiments, we conclude that 3-azido *p*HB metabolically labels PGL in *M. tb* in a highly selective manner.

**Figure 5.**
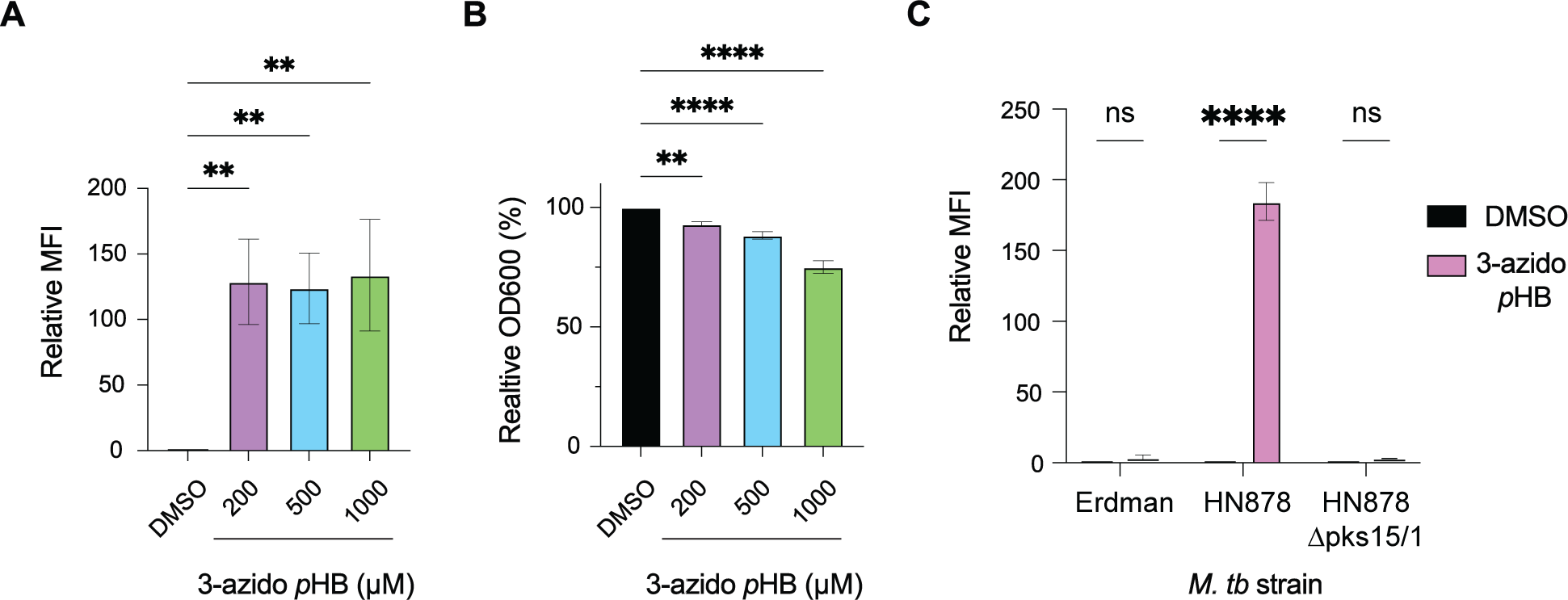
Metabolic labeling of PGL-producing *M. tb* HN878 treated with 3-azido *p*HB indicates labeling phenotype. (A) *M. tb* HN878 treated with various concentrations of 3-azido *p*HB, stained with AF488-DBCO, and analyzed by flow cytometry. Flow cytometry analysis indicates three biological replicates. Relative MFI is determined by normalizing against DMSO control. Statistical analysis was performed using a two-way ANOVA followed by a Dunnett’s multiple comparisons test. Significance is represented where \*\**p* < 0.01. (B) Effects of bacterial growth (OD600) of *M. tb* HN878 treated with various concentrations of 3-azido *p*HB. Statistical analysis was performed using a two-way ANOVA followed by a Dunnett’s multiple comparisons test. Significance is represented where \*\**p* < 0.01 and \*\*\*\**p* < 0.0001. (C) PGL-deficient *M. tb* strains (Erdman and HN878 Δpks15/1) and PGL-producing *M. tb* strain HN878 treated with 200 µM 3-azido *p*HB, stained with AF488-DBCO, and analyzed by flow cytometry. Flow cytometry analysis indicates three biological replicates. Relative MFI is determined by normalizing against DMSO control. Statistical analysis was performed using an ordinary two-way ANOVA followed by Šídák’s multiple comparisons test. Significance is represented where \*\*\*\**p* < 0.0001and ns (not significant) for *p* > 0.05.

The ability to image PGL in live mycobacteria opens the door to studies of lipid dynamics. Understanding the mobility of cell-surface lipids can inform on lipid-host interactions.^20^ In previous work, we and others have analyzed the dynamics of various mycobacterial cell wall constituents including mannosylated phosphatidylinositol,^44^ TMM,^45^ and PDIM^20^ using imaging techniques. One such technique is fluorescence recovery after photobleaching (FRAP) which is an *in vitro* confocal microscopy method to determine mobility of biomolecules.^46^ FRAP involves photobleaching a specific cellular region and monitoring fluorescence recovery over time of a fluorescently labeled biomolecule. From the rate of fluorescence recovery, the half-time of recovery (τ_1/2_) constant can be determined for a biomolecule of interest.

We therefore sought to use FRAP to investigate the membrane dynamics of PGL. We used TMM which has been previously investigated by FRAP as a comparison.^20, 45^ Mycobacteria synthesize azido-TMM upon treatment with 6-azido trehalose.^18^ We treated *M. marinum* with 3-azido *p*HB or 6-azido trehalose and stained with AF488-DBCO, to produce PGL-488 and TMM-488, respectively. In our FRAP measurements (Figure 6a and 6b), we found that TMM-488 has a half-time of recovery (τ_1/2_) of ∼10 seconds. However, PGL-488 recovered faster, with τ_1/2_ ∼3 seconds. The half-time of recovery of PGL was consistent with our previous FRAP experiment with PDIM, which had a τ_1/2_ ∼2 seconds.^20^ We then plotted the mobile fraction (Figure 6c) as determined by the plateau values from each FRAP experiment (Figure 6c). We found that the percentage of mobile lipids from TMM-488 and PGL-488 is not significantly different. From these FRAP experiments, we were able to determine that PGL is a more mobile lipid than TMM.

**Figure 6.**
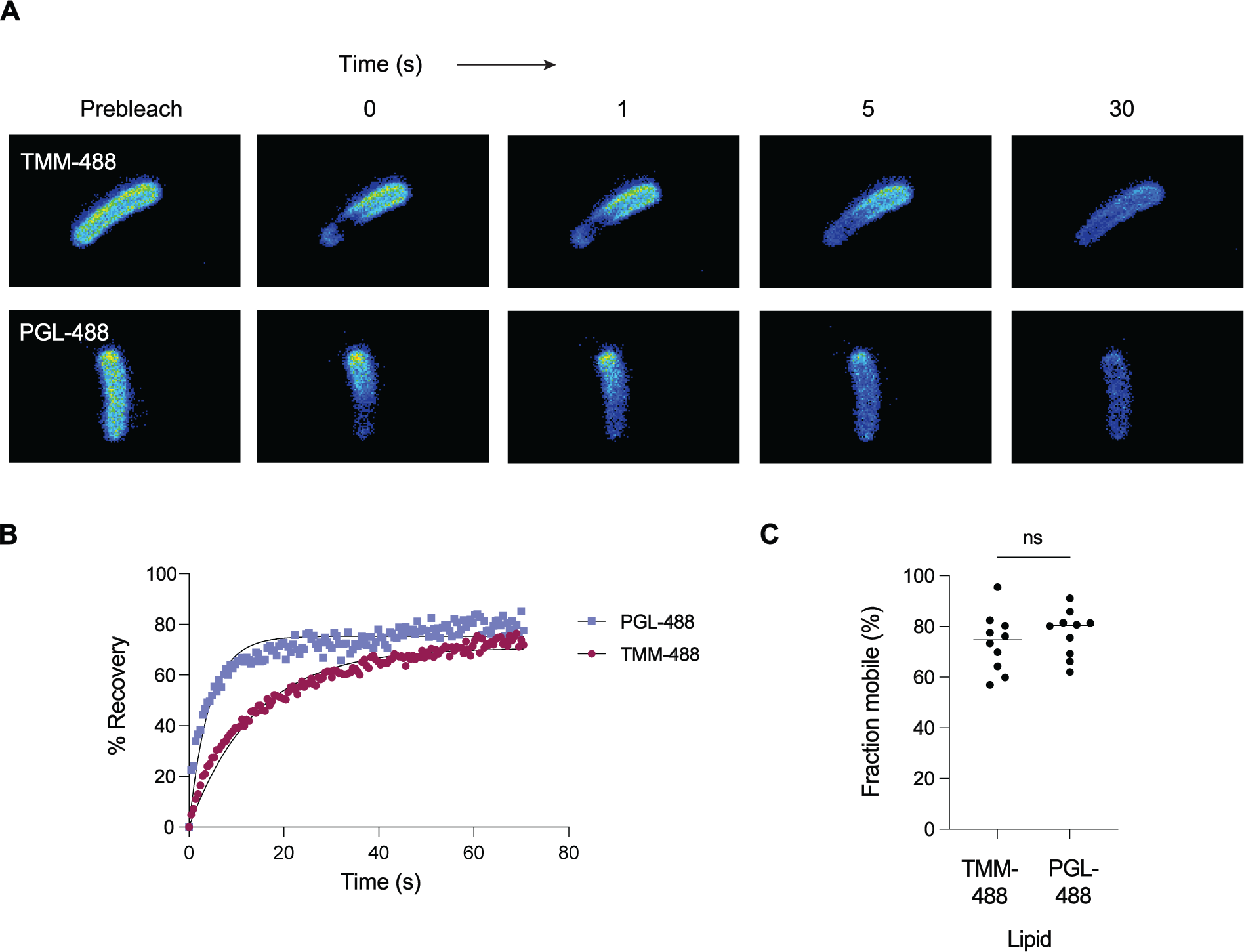
Fluorescence recovery after photobleaching (FRAP) indicates PGL has high membrane mobility in comparison to trehalose monomycolate (TMM). *M. marinum* were treated with 3-azido *p*HB or 6-azido trehalose for 18 h, stained with AF488-DBCO, and imbedded in a 1.5% agarose pad. A region of interest (ROI) was drawn around individual cells which were bleached with a 488-laser line. (A) Representative images of the fluorescence recovery after photobleaching over time. Scale bar indicates 2 µm. (B) Rates of fluorescence recovery after photobleaching. MFI values of photobleached ROIs were normalized by dividing by total fluorescence intensities of the corresponding whole cells. The plot of these values as a function of time was fitted to a non-linear regression with a one-phase association. Each symbol represents the average signal from n = 10 cells. (C) Mean mobile fraction determined as the plateau value from the fitted curves where each point represents the plateau value of an individual cell. Statistical analysis was performed using an unpaired two-tailed *t*-test represented where ns (not significant) for *p* > 0.05.

PGL is an important virulence factor hypothesized to contribute to the hypervirulence of the *M. tb* W. Lineage 2 strain HN878. Due to lack of chemical tools to tag PGL, the interrogation of PGL-host interactions during infection have been underexplored. Here, we demonstrated metabolic incorporation of a bioorthogonal handle into the mycobacterial virulence lipid PGL. We showcased the labeling with 3-azido *p*HB in the PGL-producing mycobacterial species *M. marinum* using flow cytometry and fluorescence microscopy. We identified PGL-N_3_ by mass spectrometry and determined that the 3-azido *p*HB label is selective for PGL-producing mycobacteria. Among all lipids, we determined that labeling is highly specific to PGL and established the conditions for optimally bright labeling under conditions that minimize effects on native PGL as well as other lipids. Additionally, we showed that 3-azido *p*HB metabolically labels PGL in *M. tb*. Finally, we studied PGL membrane dynamics using metabolic labeling with 3-azido *p*HB.

Studies of lipid dynamics are fundamental for understanding lipid-host interactions. We previously reported that PDIM has a high mobility and attributed its fluidity to be responsible for spreading onto host membranes during infection.^20^ Considering the structural and membrane fluidity similarities of PDIM and PGL, we hypothesize that PGL may also spread onto host membranes during infection to modulate host immunity. Although the PGL spreading mechanism has yet to be elucidated, we speculate it could be simply due to lipid shedding or by the secretion of bacterial membrane vesicles (BMVs). Indeed, PGL has been found in BMVs by performing chloroform/methanol extractions of culture supernatants.^47–48^

Future directions of this work involve visualizing the mechanisms of PGL during host infection. This involves using our metabolic labeling strategy to identify PGL distribution, trafficking, and dynamics during pathogenesis. Additionally, we theorize that our PGL metabolic labeling approach could be used as a facile method to determine if certain *M. tb* strains produce PGL without time-consuming lipid extractions and mass spectrometry analysis. We envision our metabolic labeling strategy will aid in the study of PGL during infection which may inform on therapeutic development for TB.

## Supporting information

Additional experimental details, materials, and methods, and spectra can be found in the Supporting Information.

## Acknowledgements

L.E.G would like to thank Dr. Mariko Marimoto, Dr. David Roberts, Dr. Wilson Sinclair, and Dr. Joseph Buonomo for useful discussions. L.E.G would also like to thank Dr. Manjari Mishra for technical assistance with FRAP experiments and data analysis.

## Funding sources

We are grateful for the financial support from the National Institutes of Health: R01 AI051622 (to C.R.B.), R01 AI165573 (to D.B.M.), R01 AI049313 (to D.B.M.), R01 AI158688 (to M.U.S.), and P01 AI159402 (to M.U.S). M.U.S thanks the Welch Foundation (I-1964-20210327) and the Burroughs Wellcome Fund (1017894). C.J.C was supported by the Damon Runyon Postdoctoral Fellowship. K.F.N. was supported by the NIH T32 AI007520 fellowship. Additionally, we thank the NIH High End Instrumentation grant (1 S10 OD028697-01) for the Brucker Neo-500 MHz instrument.

## Methods

### Cell lines and culture conditions

*M. marinum* was cultured at 32 °C in liquid 7H9 media containing 10% glycerol, 0.01% tween-80, and 100 µM Hygromycin B. *M. smegmatis* was cultured at 37 °C in liquid 7H9 media containing 10% glycerol, and 0.01% tween-80 shaking at 100 rpm. Starter cultures of *M. tb* strains HN878 (BEI resources), Erdman (laboratory of J. Cox, UC Berkley), and HN878 Δ*pks15/1* (a generous gift from the laboratory of C. Barry III, NIH)^8^ were grown at 37 °C in liquid 7H9 media containing 10% OADC, 1% glycerol, and 0.05% tween-80.

### Flow cytometry

*M. marinum* and *M. smegmatis* were gated based on forward and side scatter emissions excluding debris. Singlets were gated based on side scatter height vs. area. When applicable, *M. marinum* expressing wasabi were gated using the 488 nm blue laser (530/30 filter). For *M. marinum*, AF647 staining was determined by the fluorescence using the red laser (660/20 filter). *M. tb* were gated based on forward and side scatter emissions excluding debris. For *M. tb*, AF488 staining was determined by the fluorescence using the 488 nm blue laser (530/30 filter).

### General procedure for metabolic labeling experiments of *M. marinum*

*M. marinum* were cultured using 7H9 media + 0.01% tween-80 with Hygromycin B until an OD600 of 0.8-1.2. Mycobacteria were then diluted such that OD_600_ = 0.25 in 2 mL T-25 culture flasks. DMSO or azide compounds were added to the bacterial culture, with DMSO not increasing 1%. Mycobacteria were then cultured until OD600 of 0.8-1.2. The bacteria were then washed with PBS-T (3x) and once with PBS. Bacteria were then labeled with AF647-DBCO or AF680-DBCO (30 µM in PBS) for 1 h at rt in the dark. The bacteria were then washed with PBS-T (4x) and once with PBS. Bacteria were then fixed using 4% PFM and 2.5% GA for 1 h at rt in the dark. Bacteria were again washed with PBST (2x) and PBS prior to analysis by flow cytometry or microscopy.

### General procedure for lipid extractions

Large scale cultures (200 mL) of bacteria were grown until an OD600 of 0.8-1.2 shaking at 100 rpm. Bacteria were then washed 3x with PBST and once with PBS. Bacteria were then lyophilized until completely dry. Or, bacteria were then treated with a fluorophore (30 µM, 5 mL) for 1 hr at rt, washed with 4x PBST, and then 1x with PBS. Bacteria were then lyophilized until completely dry. After the cells were completely dry, the dry mass of the cells was measured. Then 20 mL of chloroform and 10 mL of methanol were added to the cells with a stir bar. The cells stirred in the organic solvent mixture overnight at rt. The cells were then filtered using Whatman 1 filter paper, rinsed with chloroform and methanol, and evaporated under reduced pressure. Lipid residue was then resuspended in chloroform and filtered using a 0.22 µm syringe filter and evaporated under reduced pressure. This filtration was repeated for a total of two times.

### General procedure for developing TLC plates

Lipid extracts were dissolved in a concentrated solution of chloroform (i.e. 20 mg/mL). A 2 µL pipette was used to apply sample to a silica gel 60 TLC plate. A small latch-lock prep TLC chamber was used to develop the TLC plates in 8:2 toluene:acetone, 95:5 chloroform:methanol, or 4:6 methanol:chloroform. After developing, TLC plates were dried with a heat gun. If fluorescent lipids were loaded onto the plate, they were scanned using a LiCor imager prior to staining with iodine or anthrone. TLC plates were then stained in a chamber containing iodine adhered to silica. After capturing an image of the TLC plate, a heat gun was used to remove most of the iodine stain. Then the TLC plate was lightly sprayed with a solution of 0.2% anthrone in H_2_SO_4_. The TLC plate was then exposed to a heat gun on high heat.

### HPLC-QTOF-MS and collision-induced dissociation (CID)-MS analysis of PGL-N_3_

Total bacterial lipids were extracted into chloroform and methanol for analysis by HPLC-MS. The lipid samples were prepared at 1 mg/ml in the starting mobile phase (50% A and 50% B), and 10 µl was injected into a reversed-phase HPLC system (Agilent 1260 series) using an Agilent Poroshell EC-C18 column (1.9-micron, 3 x 50 mm) coupled with an Agilent guard column (2.7-micron, 3 x 5 mm) and analyzed by an Agilent 6546 Accurate-Mass Q-TOF mass spectrometer. The mobile phases were A (2 mM ammonium formate in 90/10 methanol/water (v/v) and B (3 mM ammonium formate in 85/15/0.1 1-propanol/cyclohexane/water (v/v/v)). The gradients were: 0–2 min, 50% A; 2–10 min, from 50% A to 100% B; 10–15 min, 100% B; 15–17 min, from 100% B to 50% A; and 17–20 min, 50% A. CID-MS was carried-out with a collision energy of 35 V and the isolation width was set to 1.3 *m/z*.

### Metabolic labeling of *M. tuberculosis*

For each strain, starter cultures were sub-cultured into sterile square media bottles (Nalgene) at an OD_600_∼0.3 in 5mL 7H9. Cultures designated for labeling received 200-1000µM final concentration of 1 M 3-azido *p*HB or an equivalent volume of DMSO vehicle control and were wrapped in foil to protect from light. Labeled and control cultures were incubated at 37 °C shaking (100 rpm) until an OD_600_ 0.8-1.0 was reached, approximately 3 days. Following the labeling period, cultures were pelleted at 3,000xg for 3 minutes and washed 3 times with PBS+ 0.5% tween-80 and once with PBS. Bacterial pellets were then resuspended in PBS containing 10µM AFDBCO-488 or PBS alone (unstained) and incubated for 1 hour at room temperature protected from light. The bacterial suspensions were then pelleted, washed 3 times with PBS+ 0.5% tween-80, and fixed with 4% paraformaldehyde solution for 24 hours. For flow cytometry analysis of 3-azido labeling, fixed bacterial samples were washed with PBS and acquired on the BD FACSCalibur cytometer. Flow cytometry analysis was performed using FlowJo software.

### General procedure for fluorescence recovery after photobleaching (FRAP)

FRAP experiments were based on previous methods.^45^ Bacteria were cultured in the presence of 750 µM 3-azido *p*HB or 50 µM 6-azido trehalose for ∼18 h according to the general labeling procedures. The bacteria were then washed with PBS-T (3x) and once with PBS. Bacteria were then labeled with AF488-DBCO fluorophore (5 µM in PBS) for 1 h at rt in the dark. The bacteria were then washed with PBS-T (4x) and once with PBS. Low melting agarose (1.5%) pads were made, and 1 µL of bacteria were dropped onto the center of the pad. A coverslip was then put on the agar pad and the slide was sealed using nail polish. Acquisition laser power was set to 2% with bidirectional scans. Photobleaching was obtained after 4 scans using 100% bleaching power and 50 iterations. Data was obtained over 70 seconds after photobleaching. MFI values of photobleached ROIs were normalized by dividing by total fluorescence intensities of the corresponding whole cells. The plot of these values as a function of time was fitted to a non-linear regression with a one-phase association. Each symbol represents the average signal from n = 10 cells. The mean mobile fraction determined as the plateau value from the fitted curves. FRAP was performed on an inverted Zeiss 780 multiphoton laser scanning confocal microscope.

### Statistical analysis and software

GraphPad Prism 9 software was used for all statistical analyses. Significance is represented as follows: \**p* < 0.05, \*\**p* < 0.01, \*\*\**p* < 0.001, \*\*\*\**p* < 0.0001, and ns (not significant) for *p* ≥ 0.05. The specific statistical methods for individual experiments are indicated in the figure legends.

Flow cytometry data was analyzed using FlowJo 10.8.1 software. NMR data was analyzed using MNova software version 14.2.3. Mass spectrometry data was analyzed using Agilent MassHunter software. Figures were constructed using Adobe Illustrator version 26.5 software.

## Notes

### Competing Interest Statement

The authors have declared no competing interest.

### Summary of Updates

The fonts in Figure 5c aren't showing up correctly.

