## Supplemental information for "Bioorthogonal metabolic labeling of the virulence factor phenolic glycolipid in mycobacteria"

### Supporting information

Lindsay E. Guzman, C.J. Cambier, Tan-Yun Cheng, Kubra F. Naqvi, Michael U. Shiloh, D. Branch Moody, and Carolyn Bertozzi*

#### Supplementary Schemes and Figures

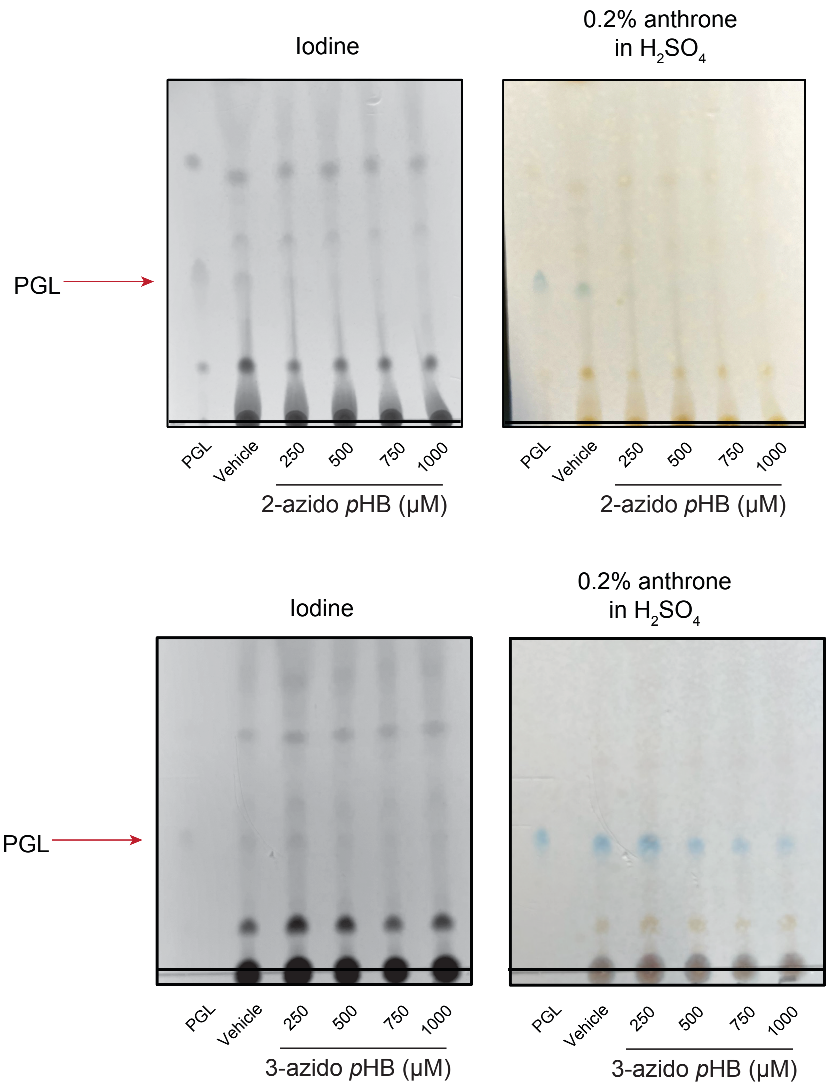


**Figure S1.** TLC plates of crude lipids extracted from *M. marinum* cultured with 2- or 3-azido *p*HB. 200 μg of crude lipids and 20 μg of purified PGL were loaded onto TLC plates and developed using 8:2 toluene:acetone. Plates were stained in 0.2% anthrone in $H_2SO_4$. The anthrone stain is used to identify compounds with sugars which will be revealed as a royal blue color. The iodine stain is a general stain used for total lipid identification.

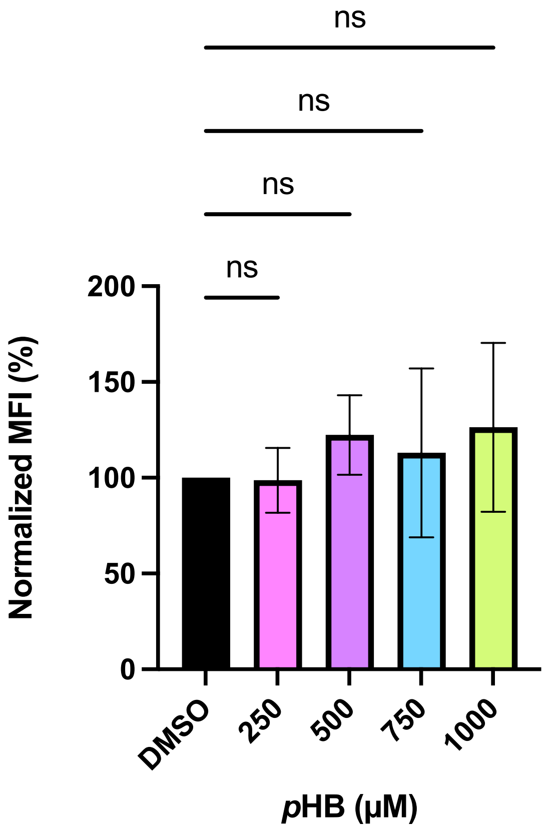


**Figure S2.** *M. marinum* cultured in various concentrations of *p*HB followed by treatment with DBCO-647 and analyzed by flow cytometry. General procedure for metabolic labeling was followed. Flow cytometry analysis represents three separate biological replicates. Relative MFI is normalized to DMSO control. Statistical analysis was performed using a one-way analysis of variance (ANOVA) followed by a Dunnett's multiple comparisons test. Significance is represented as follows: *$p \leq 0.05$, **$p < 0.01$, ***$p < 0.001$, ****$p < 0.0001$, and ns (not significant) for $p > 0.05$.

Supplementary Table 1

| PGL | | | | PGL-N3 | | | | |
|---|---|---|---|---|---|---|---|---|
| neutral molecule formula | [M+NH4]+ calculated m/z | [M+NH4]+ detected m/z | intensity (area) | neutral molecule formula | [M+NH4]+ calculated m/z | [M+NH4]+ detected m/z | mass error ppm | intensity (area) |
| C94H174O10 | 1481.3445 | 1481.344 | 1,038,938±110,812 | C94H173N3O10 | 1522.3459 | not detected | | |
| C95H176O10 | 1495.3602 | 1495.359 | 712,617±48,059 | C95H175N3O10 | 1536.3616 | 1536.365 | 2.3 | 66,001±645 |
| C95H178O10 | 1497.3758 | 1497.375 | 304,183±19,881 | C95H177N3O10 | 1538.3772 | not detected | | |
| C96H178O10 | 1509.3758 | 1509.376 | 1,548,098±124,234 | C96H177N3O10 | 1550.3772 | not detected | | |
| C96H180O10 | 1511.3915 | 1511.390 | 337,981±11,557 | C96H179N3O10 | 1552.3929 | 1552.403 | 6.8 | 49,841±5,499 |
| C97H180O10 | 1523.3915 | 1523.393 | 4,417,171±278,616 | C97H179N3O10 | 1564.3929 | 1564.394 | 0.6 | 206,331±3,679 |
| C97H182O10 | 1525.4071 | 1525.407 | 753,652±63,171 | C97H181N3O10 | 1566.4085 | not detected | | |
| C98H182O10 | 1537.4071 | 1537.407 | 1,719,998±124,695 | C98H181N3O10 | 1578.4085 | not detected | | |
| C98H184O10 | 1539.4228 | 1539.424 | 2,776,249±166,514 | C98H183N3O10 | 1580.4242 | 1580.424 | 0.1 | 160,966±8,394 |
| C99H184O10 | 1551.4228 | 1551.421 | 4,578,012±364,026 | C99H183N3O10 | 1592.4242 | 1592.435 | 6.8 | 36,801±6,232 |
| C99H186O10 | 1553.4384 | 1553.436 | 952,323±71,916 | C99H185N3O10 | 1594.4398 | not detected | | |
| C100H186O10 | 1565.4384 | 1565.437 | 1,123,848±71,813 | C100H185N3O10 | 1606.4398 | 1606.449 | 5.8 | 20,635±2,192 |
| C100H188O10 | 1567.4541 | 1567.456 | 4,640,701±309,886 | C100H187N3O10 | 1608.4555 | 1608.452 | 2.2 | 29,151±5,547 |
| C101H188O10 | 1579.4541 | 1579.455 | 3,088,382±163,416 | C101H187N3O10 | 1620.4555 | not detected | | |
| C101H190O10 | 1581.4697 | 1581.464 | 315,240±16,810 | C101H189N3O10 | 1622.4711 | not detected | | |
| C102H190O10 | 1593.4697 | 1593.469 | 496,021±55,504 | C102H189N3O10 | 1634.4711 | not detected | | |
| C102H192O10 | 1595.4854 | 1595.486 | 1,509,159±100,619 | C102H191N3O10 | 1636.4868 | not detected | | |

**Supplemental Table 1.** HPLC-MS positive mode analysis of PGL and PGL-$N_3$ from the extracted lipids of *M. marinum* grown in media supplemented with 750 μM 3-azido *p*HB. The known PGLs were detected as ammonium adducts, $[M + NH_4]^+$. For PGL-$N_3$, the theoretical molecular formula and mass were deduced by replacement of a proton for $N_3$. Seven PGL-$N_3$ species were identified by near coelution with PGL and matching the calculated masses to predict masses with errors less than 10 ppm.

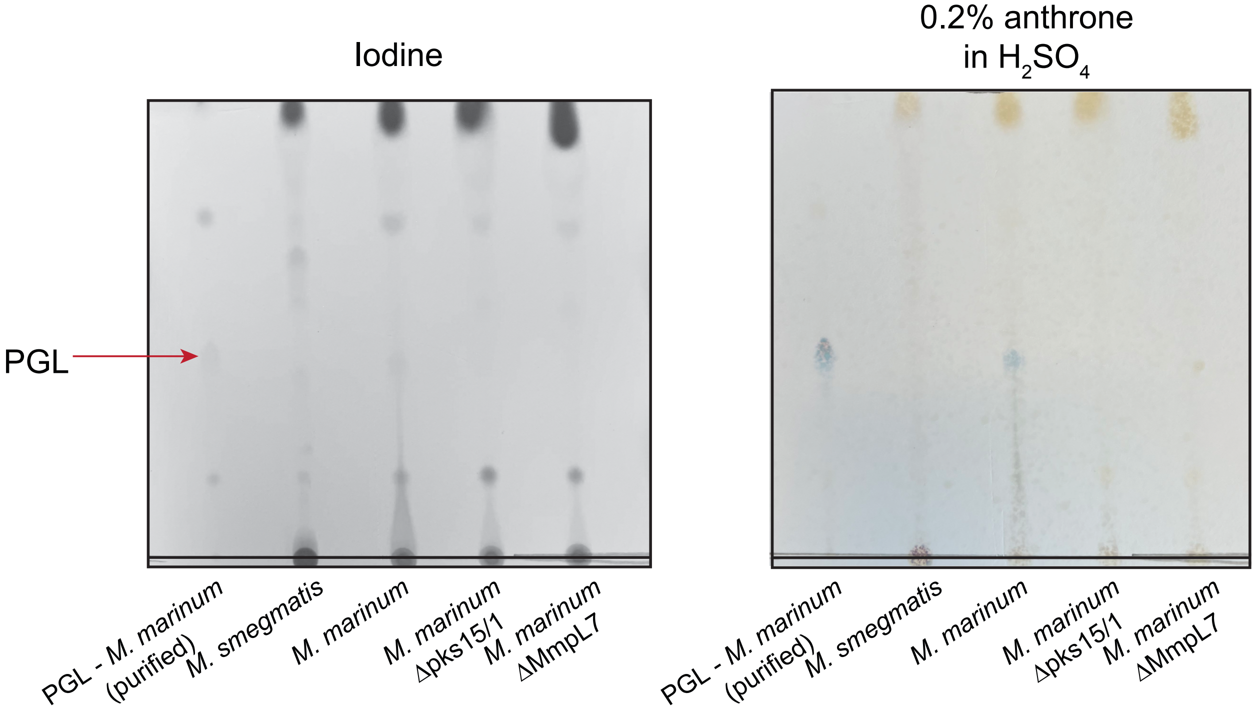


**Figure S3.** Crude lipid extracts from PGL-deficient *M. smegmatis*, PGL-producing WT *M. marinum*, and PGL-deficient *M. marinum* mutants indicate presence or absence of PGL. Purified PGL from M. marinum was used as a standard. Crude lipid extracts (100 µg) and purified PGL- *M. marinum* (5 µg) were loaded onto a silica gel 60 TLC plate and developed in 8:2 toluene:acetone. Compounds with sugars are visualized as a blue color in the anthrone stain. The iodine stain is a general stain used to visualize total lipids.

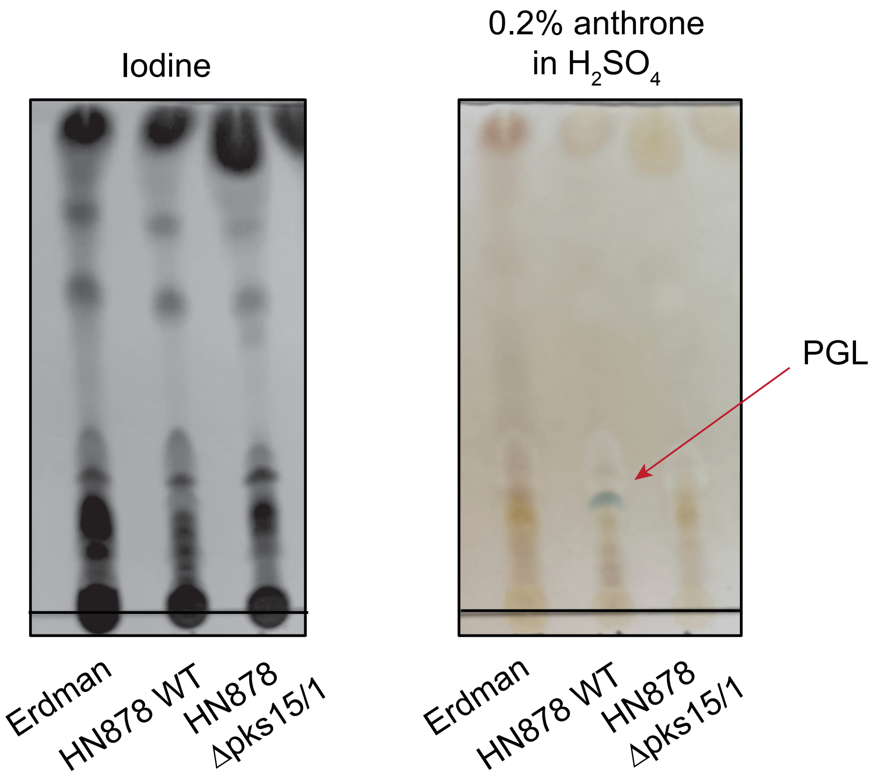


**Figure S4.** Crude lipid extracts from *M. tb* strains Erdman, HN878 WT, and HN878 Δpks15/1. Crude lipid extracts (100 µg) were loaded onto a silica gel 60 TLC plate and developed in 95:5 chloroform:methanol. Compounds with sugars are revealed as a blue color in the anthrone stain. The iodine stain is a general stain used to check for total lipids.

#### Supplementary methods

**General methods for synthesis**. Materials and reagents were obtained from commercial sources without further purification unless otherwise noted. Analytical TLC was performed on Millipore Sigma glass-backed Silica gel 60 F254 plates. Prep TLC was performed on Millipore Sigma glass-backed 2 mm thick Silica gel 60 $F_{254}$ plates with concentration zone. TLC was analyzed using iodine adhered to silica and 0.2% anthrone in $H_2SO_4$ stain. Column chromatography was performed using normal-phase silica gel 60. $^1$H and $^{13}$C NMR spectra was obtained using a Brucker Neo-500 MHz instrument with chemical shifts in ppm ($\delta$) referenced to solvent peaks. Flow cytometry analysis of *M. marinum* was performed on a Novocyte Penteon flow cytometer at the Stanford shared FACS facility. Flow cytometry analysis of *M. tuberculosis* BD FACSCalibur cytometer at UT Southwestern. High resolution mass spectrometry characterization of azide compounds was performed on an Thermo Orbitrap Fusion nano LC/MS instrument at the Stanford Mass Spectrometry facility. FRAP was performed at the Stanford microscopy facility on an inverted Zeiss 780 multiphoton laser scanning confocal microscope. Lipidomics analysis of crude extracts was performed on a reversed-phase Agilent 1260 series HPLC system and an Agilent 6546 Accurate-Mass Q-TOF mass spectrometer.

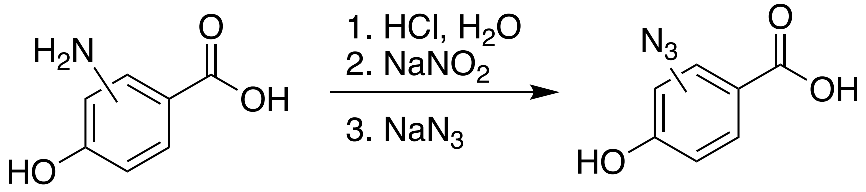


**General procedure for synthesis of 3- and 2-azido *p*HB**. *WARNING: the synthesis of azide compounds is dangerous and can result in explosions and/or production of toxic gases. It is recommended to synthesize these molecules at small scale (i.e. 1 mmol) behind a blast shield. Proceed with caution.*

Aniline (2- or 3-amino pHB) starting materials (0.150 g, 1 mmol) were resuspended in 2.5 mL of water in a round bottom flask. The round bottom flask was submerged in an ice bath and conc. HCl (0.25 mL) was added slowly. The flask stirred at 4 °C for 10 minutes. Next, $NaNO_2$ (0.076 g, 1 mmol) was added to the flask and stirred at 4 °C for 10 minutes.Then $NaN_3$ (0.085 g, 1.3 mmol) was added to the flask and stirred at 4 °C for and additional 30 minutes. After 30 minutes the flask was removed from the ice bath and warmed to rt. Reaction was monitored using 1:1 acetone:hexanes. After ~2 hrs when the starting material was consumed, an additional 10 mL of water was added to the flask and the product was extracted with 3 x 10 mL of ethyl acetate. The organic layer was then washed with brine and dried using magnesium sulfate. A short plug-like silica column was packed in hexanes and the crude product was purified using a gradient solvent system of starting with hexanes and ending with 1:1 acetone:hexanes to achieve the azido product as an off-white solid. Yield of 2- and 3-azido *p*HB was ~70%.

**3-azido *p*HB**: $^{1}$H NMR (500 MHz, d$^{6}$-DMSO): δ 7.61 (dd, *J* = 2.2, 10 Hz, 1 H), 7.4 (d, *J* = 2.2 Hz, 1 H), 6.94 (d, *J* = 9.9, 1 H). $^{13}$C NMR (500 MHz, d$^{6}$ DMSO): 166.52, 154.53, 127.82, 125.78, 122.32, 121.94, 116.07. HR ESI MS: calculated for C7H5N3O3[-H]$^{-}$ m/z, 178.0247; found 178.0255.

**2-azido *p*HB**: $^{1}$H NMR (500 MHz, d$^{6}$-DMSO): δ 10.52 (s, 1H), 7.71 (d, J = 5, 1H), 6.64 (m, 2H). $^{13}$C NMR (500 MHz, d$^{6}$-DMSO): δ 165.75, 161.77, 141.30, 133.88, 114.00, 112.44, 107.09. HR ESI MS: calculated for C7H5N3O3[-H]$^{-}$ *m/z*, 178.0258; found 1278.0259.

#### NMR Spectra

**3-azido *p*HB**: $^{1}$H NMR ($d^6$-DMSO, 500 MHz)

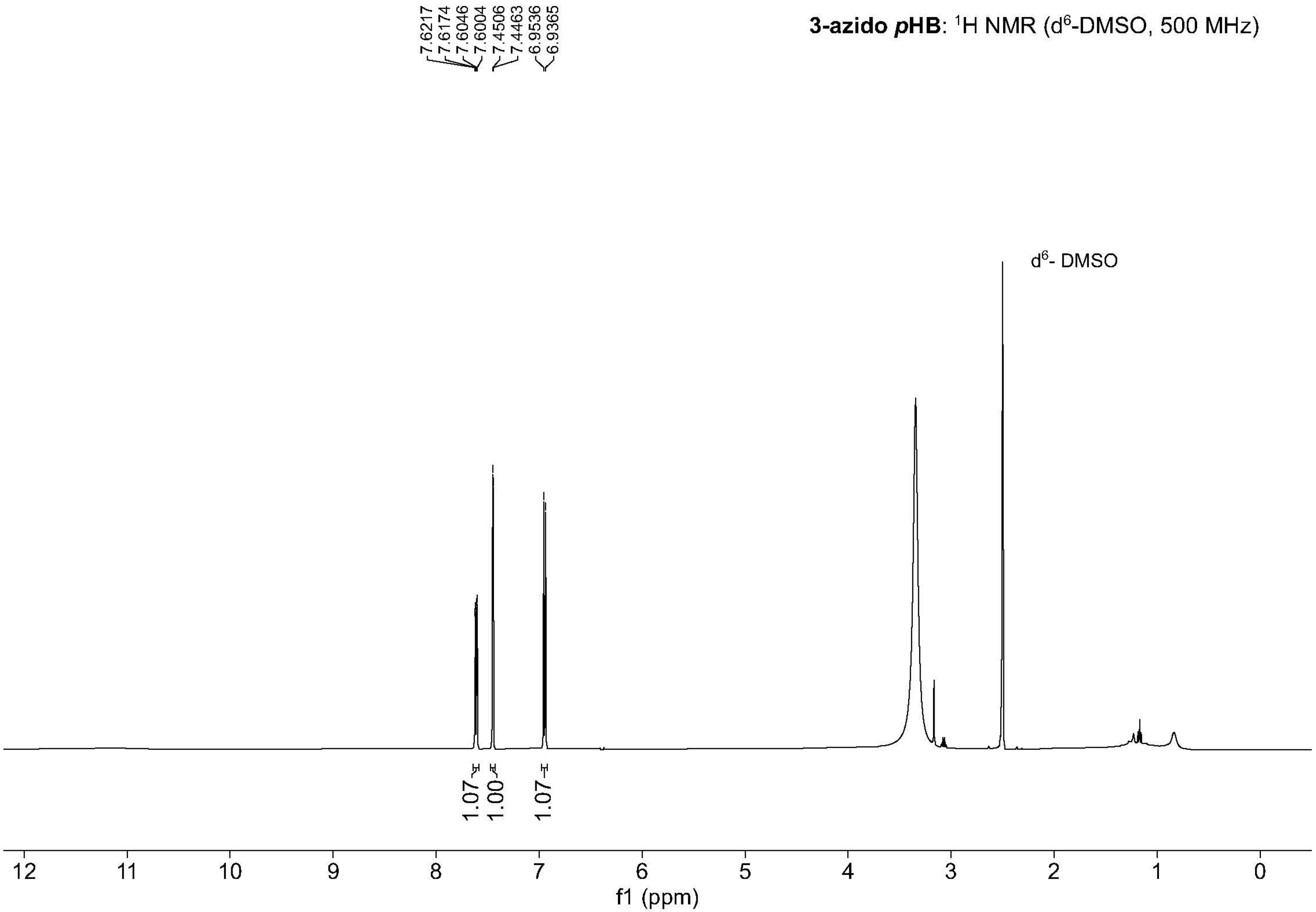

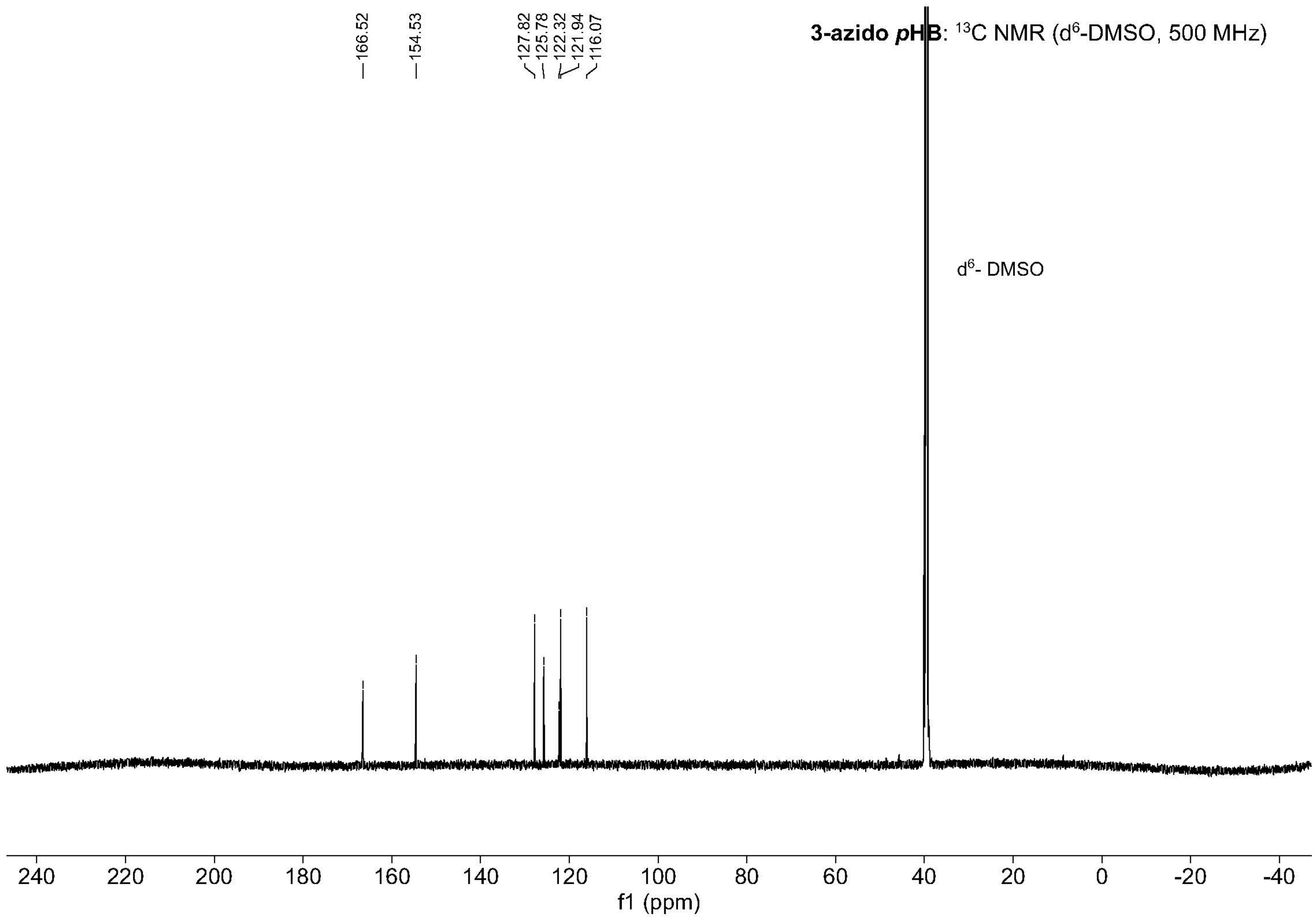
3-azido pHB: 13C NMR (d6-DMSO, 500 MHz)
166.52
154.53
127.82
125.78
122.32
121.94
116.07
d6- DMSO
240 220 200 180 160 140 120 100 80 60 40 20 0 -20 -40
f1 (ppm)

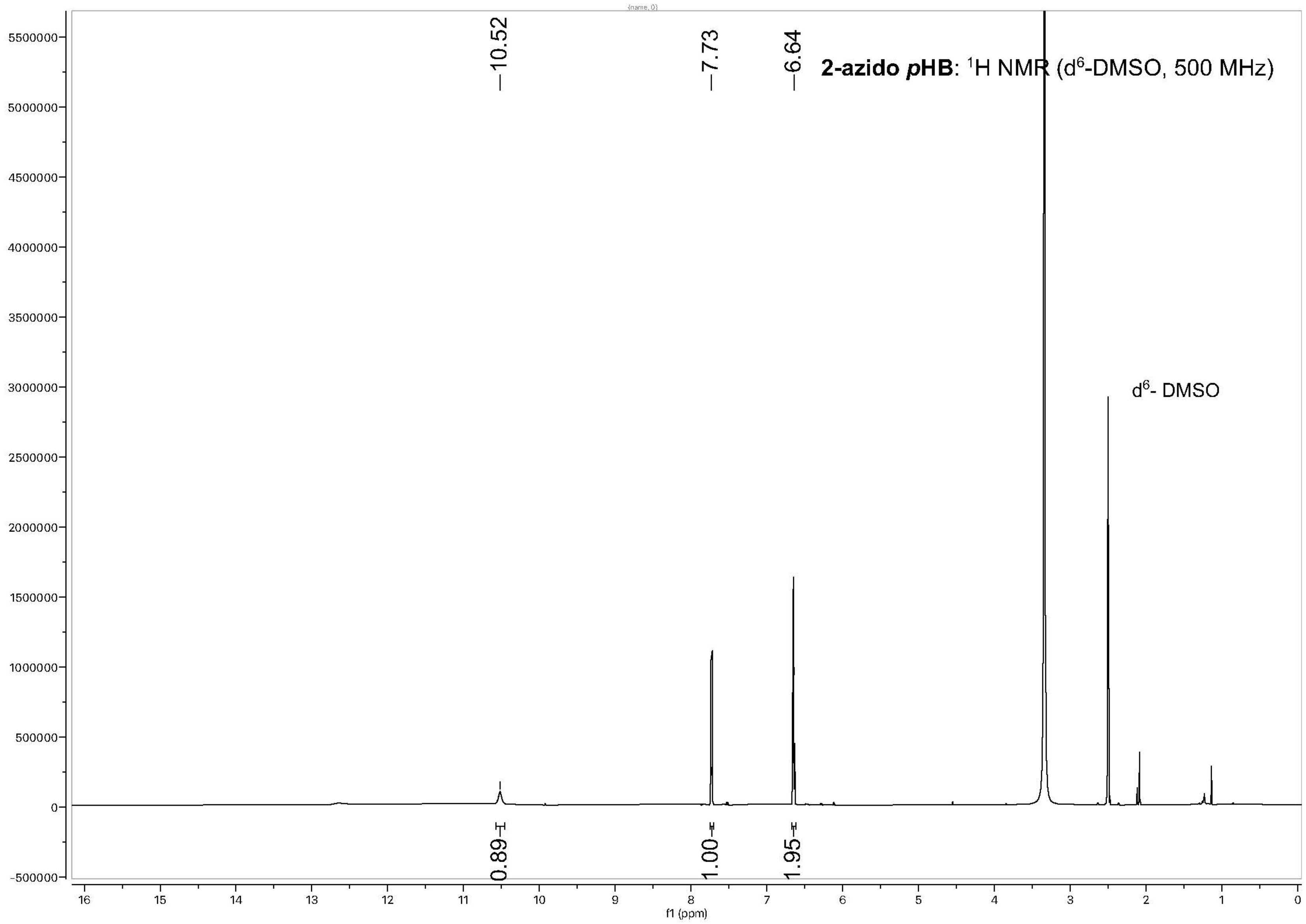
2-azido pHB: 1H NMR (d6-DMSO, 500 MHz)
10.52
7.73
6.64
d6- DMSO
0.89
1.00
1.95
f1 (ppm)

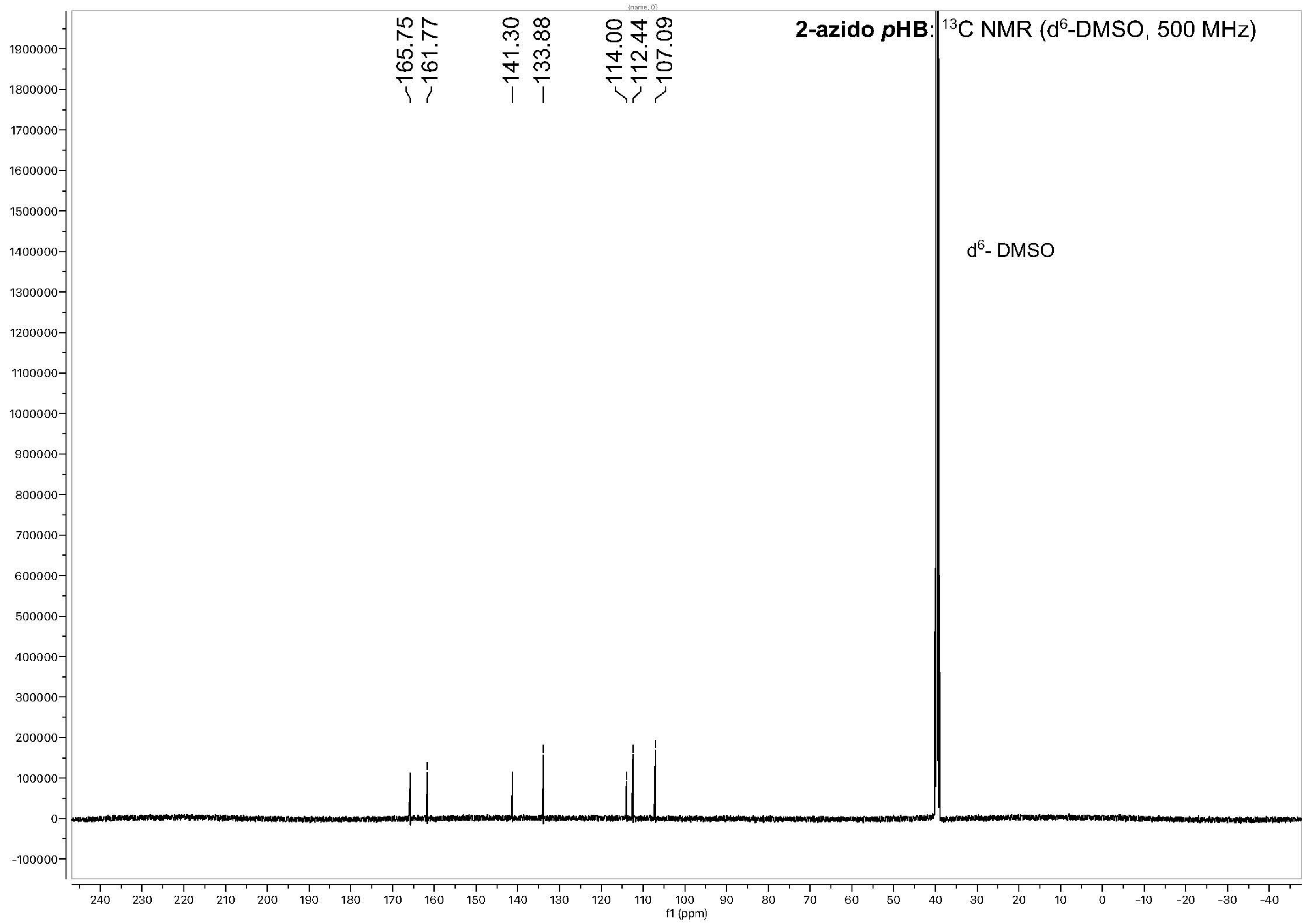

2-azido pHB: 13C NMR (d6-DMSO, 500 MHz)
165.75
161.77
141.30
133.88
114.00
112.44
107.09
d6- DMSO
f1 (ppm)
